# Electrochemical guided mode resonance biosensor for simultaneous refractive index and electrochemical measurements

**DOI:** 10.1101/2025.07.16.665075

**Authors:** Stephen Thorpe, Sam Blair, Giampaolo Pitruzello, Casper Kunstmann, Christopher P Reardon, Thomas F Krauss, Steven D Johnson

## Abstract

A comprehensive miniaturised biosensing platform needs to detect multiple analytes, which often rely on different transduction mechanisms for their detection and quantification. This necessitates the development of multimodal sensors capable of simultaneous measurements without interference between measurement modalities. Guided mode resonance (GMR) biosensors are highly sensitive to refractive index changes and well suited to out-of-plane optical coupling, but have not previously been implemented in a multimodal configuration. Here, we present the electrochemical guided mode resonance (EC-GMR) sensor, a Si_3_N_4_ GMR photonic grating integrated with an indium tin oxide (ITO) electrode that enables concurrent optical and electrochemical sensing. Finite difference time domain (FDTD) simulations show that the resonance wavelength of the GMR shifts as a function of ITO thickness and applied bias. These findings are validated experimentally using EC-GMR devices fabricated with a range of ITO thicknesses. We demonstrate that the EC-GMR achieves a refractive index sensitivity of 84.4 nm/RIU while simultaneously measuring the double-layer capacitance. To highlight its multimodal capability, we characterise the optical and electrochemical responses of methylene blue, a widely used redox reporter in biosensing. By integrating photonic and electrochemical modalities in a compact format, the EC-GMR platform provides critical analytical information beyond that which could be obtained from a traditional single-modality sensor. This approach extends the analytical utility of GMR sensors and represents a robust, adaptable solution for diverse biosensing applications, from clinical diagnostics to environmental monitoring.

## Introduction

A comprehensive understanding of biological systems for diagnostic and research purposes requires untangling a system of complex molecular interactions. The majority of biosensing techniques are “single domain”, relying on a single method of transduction such as optical, electrochemical or mechanical, limiting the total information that can be extracted and the range of biological interactions that can be investigated^1–3^. Multi-domain sensors combine multiple transduction techniques to simultaneously probe different elements of a biological interaction increasing the amount of information obtained compared to single domain biosensors ^4^. A recent area of interest in multimodal sensing has been the combination of electrochemical and photonics biosensors. The integration of these modalities allows simultaneous measurement of refractive index and charge transport to extend the range of the interactions that can be detected and the richness of analytical information that can be obtained. For example, a sensor that combines photonic and electrochemical transduction can interrogate both protein ligand interactions through the associated change in refractive index, alongside electrical characterisation of surface immobilised enzyme activity that cannot be measured with a single domain sensor alone for example in antibiotic binding and enzymatic turnover of β-lactam antibiotics by β-lactamase enzymes ^5,6^. Moreover, the integration of the electrochemical and the photonic domains not only allows us to combine information contained within the two, complementary measurements, but also to actively perform electrochemistry simultaneous with optical characterisation, for example to provide insight into electrochemical processes such as the role of redox active signalling in *Pseudomonas aeruginosa* biofilm formation or to permit electrochemically driven surface modification^7^.

The most established electrochemical and optical multimodal technique is Electrochemical Surface Plasmon Resonance (EC-SPR). EC-SPR uses the Au surface of the SPR sensor as the working electrode and integrates standard counter and reference electrodes for performing electrochemical measurements^8^. The multimodality of the EC-SPR has been used for the parallel detection of glucose and IgG using a surface functionalised with a mixed layer of glucose oxidase and anti-IgG, where the glucose oxidase activity was monitored electrochemically and the binding of IgG detected optically through the SPR angle shift ^9^. Electrical control of the surface charge afforded by EC-SPR can also be exploited to improve the performance of SPR detection compared to single modality devices ^10,11^. EC-SPR also provides additional value over single modality sensors when applied to electron transport processes that regulate cell growth and viability of both prokaryotes and eukaryotes ^12,13^.

In contrast to the metallic materials used in plasmonic sensors such as SPR, photonic biosensors are typically fabricated in loss-less dielectric materials. It is thus necessary to integrate a conductive material such as transparent conductive oxides or conductive polymers into the structure to add the electrochemical modality. This approach to combining electrochemical and photonic modalities is employed in electrochemical optical waveguide lightmode spectroscopy (EC-OWLS) and Fibre based lossy mode sensors (LMR). EC-OWLS has been used to characterise the electric-field mediated surface-adsorption of proteins and charged polymers such as poly-L-lysine, and to probe the activity of surface immobilised redox-active molecules such as cytochrome c ^14–17,^. EC-OWLS however requires high-quality He-Ne lasers, precise control over the angle of incidence, and facet out-coupling of the detected signal, making them less suitable for cost-effective implementations and environments with high levels of mechanical or optical noise. Electrochemical LMR has been used to characterise electropolymerisation of isatin, a drug precursor compound, using a combination of cyclic voltammetry and optical measurements of the change in refractive index^18,19^. The issue with LMR is that its optical response under applied bias voltage has been shown to be dependent on both the properties of the cladding and of the surrounding electrolyte ^19^, so it is difficult to extract photonic and electrochemical information independently.

We have previously demonstrated an electrophotonic silicon biosensor using a doped optical ring resonator that supports both optical sensing and electrochemical measurements^20^. In a DNA hybridisation assay, only the combined optical and electrochemical signals confirmed the conformational change of a redox-labelled DNA hairpin. Optical detection alone lacked sensitivity, while the electrochemical signal was not uniquely attributable to the structural change. ^20^These exemplars clearly demonstrate the value of simultaneous electrochemical and optical measurements, yet several challenges remain preventing their wider adoption. First, electrically induced resonance shifts in EC-SPR and LMR sensors complicate parallel electrochemical and refractive index measurements, as the shifts cannot be attributed solely to refractive index changes during voltametric scanning^21^. Although this effect can optically track electrochemical processes, true multimodal integration requires the signals to be independent^22^. Second, quantitative validation of both refractive index sensitivity and electrochemical response, across faradaic and non-faradaic system.is needed to fully demonstrate the technique’s potential, which, to our knowledge, has not yet been achieved. Finally, a key technological limitation with existing multimodal biosensing technology such as EC-OWLS is the requirement for complex and sensitive measurement apparatus for both sensing modalities, which often precludes their use for applications where portability or mechanical robustness are critical.

We recently demonstrated a biosensor platform based on photonic guided mode resonances (GMR) which offer the same evanescent wave sensing principle and performance of SPR without the need for complex instrumentation. Moreover, the entire device (GMR sensor, optical system and control electronics) can be implemented in a small and robust, instrument reducing sensitivity to mechanical vibrations such as those in industrial manufacturing systems^23,24^. Electrically tunable filters using GMR surfaces have been previously demonstrated but there has been no work to date on combining optical and electrical sensing modalities using a GMR biosensor ^25^.When combined with low cost potentiostat electronics, an electrochemical GMR sensor has the potential to be a highly robust and economical multimodal sensing platform ^26^.

A GMR sensor is a 1D periodic sub wavelength grating, which acts as both a diffraction grating and a waveguide. The guided mode is scattered at the grating interfaces and by tuning the grating parameters including thickness, period and filling fraction, deconstructive interference between the 0^th^ order and guided mode results in a high reflectance peak at the resonance wavelength ^27^. The resonance wavelength is dependent on the local refractive index (RI) within 200 nm of the sensor surface. When functionalised with selective biomolecules, such as antibodies, the change in RI at the GMR surface that occurs following formation of the antibody–antigen complex is detected by the corresponding change in resonance wavelength^28^. GMR sensors have also been used for cell sensing such as bacterial biofilm formation and adherence in response to antibiotics and spatially resolved signalling protein secretion in hepatocyte cells^29,30^.

Here we present electrochemical GMR (EC-GMR), a multimodal GMR sensor with an integrated ITO working electrode that can be used for parallel optical and electrochemical sensing (**Fig. 1**). We explore theoretically the influence of an ITO layer on the GMR resonance and electric field distribution for the resonant mode and how modulation of the accumulation layer refractive index at the ITO media interface affects the E-GMR resonance. We then fabricated the proposed structure, in order to experimentally determine the relationship between optical resonance and applied bias voltage. Finally, we demonstrate the capabilities of the multimodal EC-GMR sensor by performing refractive index sensing in parallel with electrochemical measurements. In particular, we show that the sensor is capable of supporting quantitative measurements of both Faradaic and non-Faradaic electrochemical processes alongside parallel quantitative measurements of local refractive index which, when combined, provide critical analytical information beyond what is possible form a single-modality. Together, our data lay the foundation for future applications of this combined optical-electrochemical sensor in basic and applied biochemical research.

**Figure 1.**
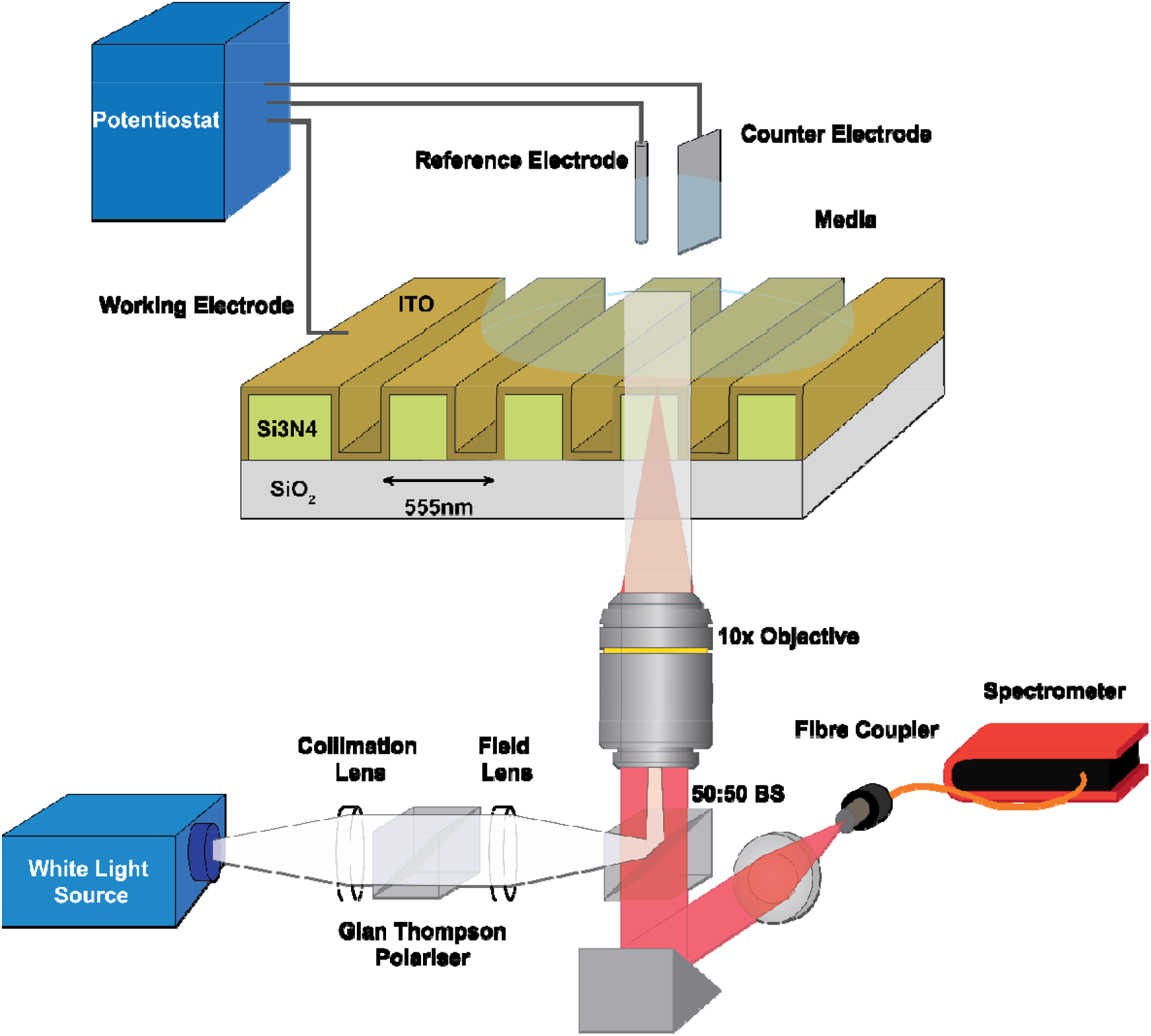
Schematic of the EC-GMR and measurement setup. The EC-GMR consists of a 555nm period GMR 150 nm deep Si_3_N_4_ grating on glass coating in a layer of ITO. For measuring the resonance wavelength a broadband white light source is collimated and the polarisation setup by a Glan Thompson polariser mounted in a rotation mount. The incident light is focused into the back focal plane of the 10x objective by the field lens producing a collimated incident beam at the EC-GMR. The reflected light from the EC-GMR is fibre coupled into the spectrometer. For electrochemical measurement the ITO surface is used as the working electrode, with the Ag/AgCl reference electrode and Pt counter electrode placed in solution above the EC-GMR.

## Results and Discussion

### Simulation of a functional hybrid ITO Si_3_N_4_ EC-GMR biosensor

To verify that a GMR structure can still support a resonance when coated with a uniform and continuous film of ITO, we performed finite difference time domain (FDTD) simulations on a GMR structure consisting of a 150 nm thick Si_3_N_4_ grating with 555 nm grating period and 80% filling fraction. The glass substrate and media layers were assumed to be infinite. Firstly, we simulated the GMR structure with increasing thicknesses of ITO from 0 – 100 nm. To account for a potentially non-conformal coating of the ITO, we varied the ITO thickness on the sidewalls of the GMR trenches following deposition. The sidewall thickness was varied from 10 - 50% of the thickness deposited on the top of the grating.

We observe that the structure is theoretically capable of supporting the TE mode of the resonance for all thicknesses of ITO (**Fig 2 a**) and displays the asymmetry of the Fano line shape due to coupling of the Bragg reflectance and the Fabry Perot thin film resonance ^31^ As the ITO thickness increases, the resonance wavelength for the TE mode increases from 810 nm at 0 nm ITO thickness to 890 nm for a 100 nm thick ITO layer. The Q factor also theoretically increases with ITO thickness due to increased confinement of the electric field within the grating structure. The electric field intensity in an ITO coated grating is confined within the centre of the structure and in the trenches of the grating, showing a strong overlap with the ITO in the centre (**Fig. 2b**). Increasing the sidewall coverage of the ITO further increases the resonance wavelength for all thicknesses, as expected by increasing the effective refractive index experienced by the resonant mode (**Fig supplementary 1)**.

**Figure 2.**
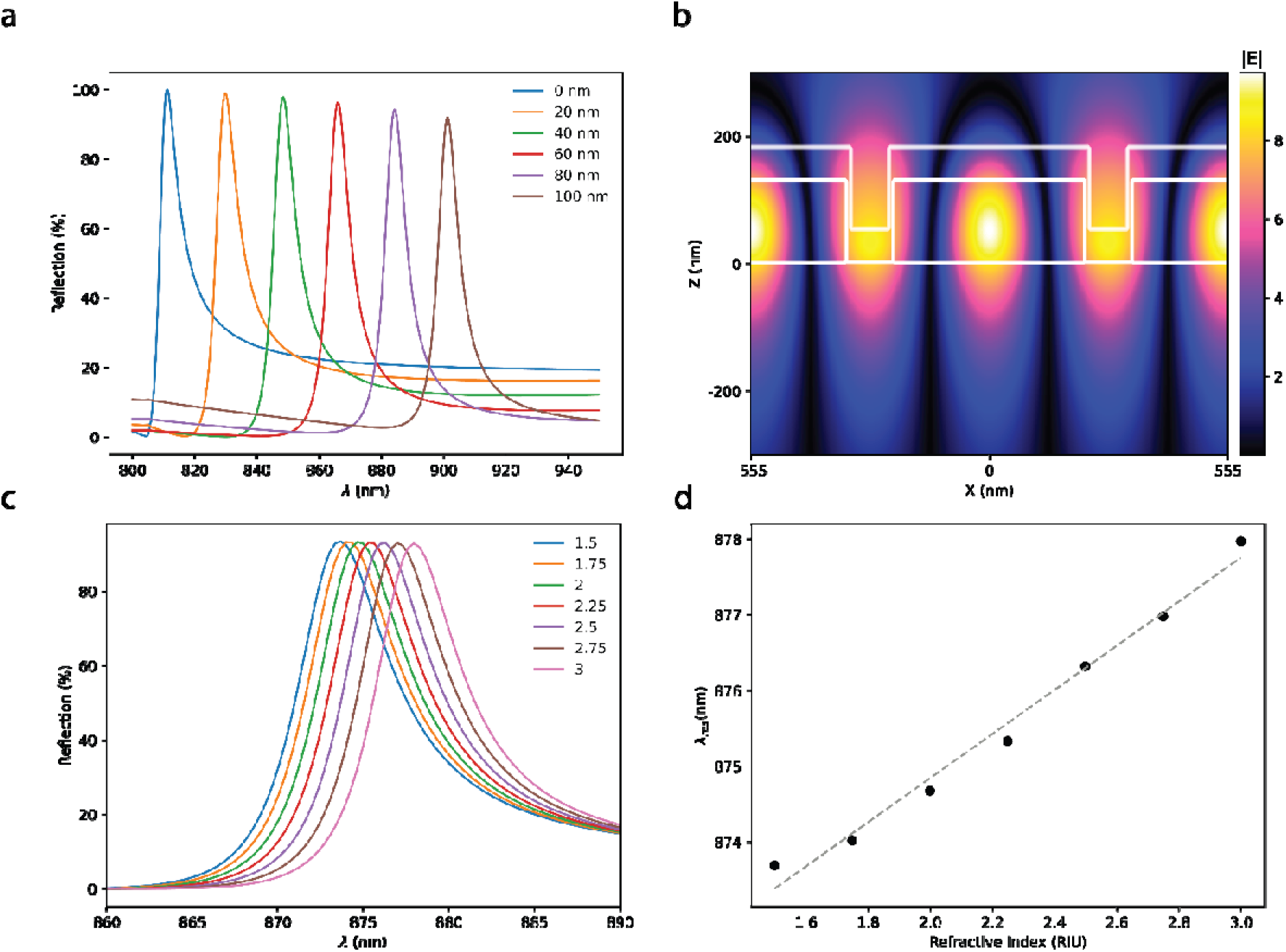
Simulation of EC-GMR resonance wavelength with ITO thickness a Simulated. TE reflectance spectra of EC-GMR with different thicknesses of ITO from 0 to 100 nm. **b** Electric field distribution of the TE mode of the EC-GMR coated with 60nm of ITO. **c** Simulated TE reflectance spectra of EC-GMR for increasing refractive indexes of a 1nm accumulation region at the ITO media interface. **d** Peak resonance wavelength against accumulation refractive index change, R^2^ = 0.98.

To investigate how the resonance changes due to electrically induced electron accumulation/depletion within the ITO layer, we simulated a 60 nm thick layer ITO with 40% infilling of the trenches and assuming a 1 nm thick accumulation region at the interface between the ITO and surrounding media. Here, the RI of the accumulation layer was changed (from 1.5 to 3 RIU) to simulate the shift in RI due to applied bias (**Fig. 2c**). The resonance is still supported as expected and the peak resonance wavelength, λ_res_, increases linearly with refractive index of the accumulation layer at 2.9 nm/RIU (**Fig. 2d**). This suggests that calibration of the electrically induced resonance would be required for parallel optical and electrochemical measurements. It is important to note the overlap between the ITO accumulation layer and the resonant electric field is much greater for the TE mode compared to the TM mode, which will contribute to a greater resonance wavelength change (**Fig supplementary 2**). FDTD simulations demonstrate that the resonant mode within the GMR is robust to the addition of an optical lossy high refractive index material and therefore viable as a multimodal biosensing platform.

### EC-GMR Fabrication

The grating structure was fabricated using electron beam lithography and RIE to etch the grating into a 150 nm thick Si_3_N_4_ layer deposited onto a glass substrate. ITO is then deposited on top of the grating by electron beam evaporation. Electron micrographs of an ITO-coated GMR (**Fig. 3a**) reveal ITO deposition on the top of the grating and on the sidewalls. Moreover, the ITO within the grooves of the grating has greater thickness and surface roughness than that on the top of grating. All of the samples fabricated showed a single resonance peak in the 800 - 900nm region when illuminated with a broadband white light source (**Fig. 3b**). The resonance wavelength increases with ITO thickness as predicted by simulations (**Fig. 2a**) due to the increase of the effective index of the structure. The EC-GMR is theoretically capable of supporting the TM mode but it was not observed in any of the structures that were measured.

**Figure 3.**
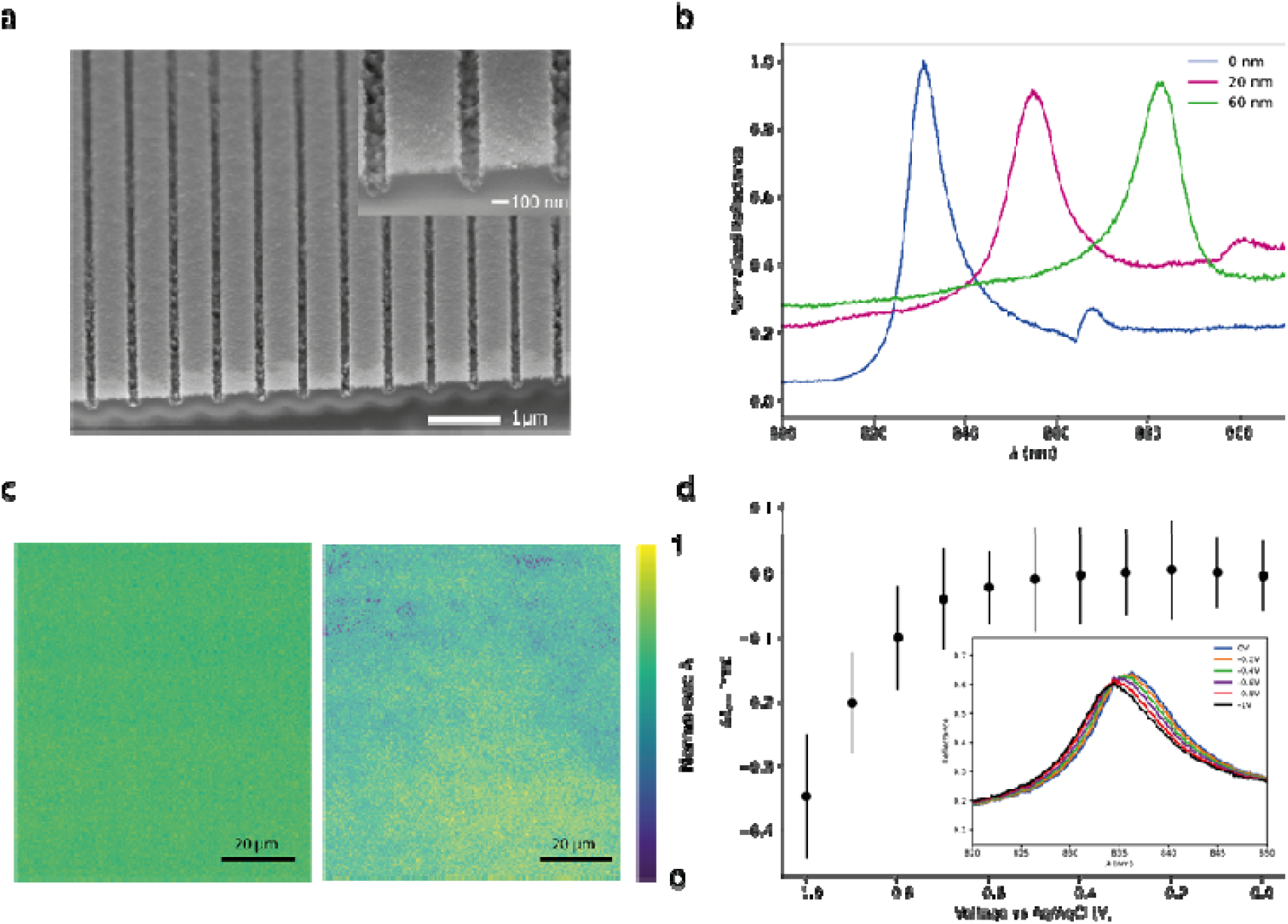
EC-GMR Grating Structure and Resonance Modulation by Bias Voltage: **a** SEM image of EC-GMR grating which has been cleaved through to expose the profile of the grating. ITO thickness is 62 nm. inset SEM image of EC-GMR grating grooves. **b** Reflectance spectra of the TE mode for the Si3N4 grating, 20 nm ITO EC-GMR grating and 60 nm EC-GMR grating. Reflectance has been normalised to an Ag mirror. **c** Hyperspectral image of resonance wavelengths for Si3N4 GMR grating (left) and EC-GMR with 60 nm thick ITO layer (right). The resonance wavelength has been normalised between 0 and 1 for both images. Image area is 181×181 µm. **d** Change in resonance wavelength Δλ_res_ for 60 nm EC-GMR in 100 mM KPi buffer with increasing bias voltage from 0 to −1. Reflectance spectra shown in inset.

Simulations also predicted an increase in Q factor with increasing ITO thickness, as the propagation length of the mode will be reduced by the optical losses introduced by the ITO. In contrast, experimentally we observed a broadening of the resonance with increasing thickness of ITO and a subsequent decrease in Q factor compared to the uncoated Si_3_N_4_ grating (0 nm thick ITO in Fig. 3b).

This is most likely caused by light scattering due to the inhomogeneity of the ITO film which introduces many additional randomly orientated surfaces from which the guided mode can scatter.

Hyperspectral imaging of an uncoated Si_3_N_4_ grating shows that the resonance wavelength is consistent across the entire grating, with a standard deviation of only 1.02 nm in resonance wavelength (**Fig. 3c left**). In contrast, the standard deviation of the resonance wavelength in the ITO coated EC-GMR is 1.54 nm, over 50% greater than the Si_3_N_4_ grating. Moreover, the resonance is inhomogeneous across the grating, with areas of high and low resonance contributing to the overall broadening of the reflectance peak, again likely associated with inhomogeneities within the evaporated ITO film (**Fig. 3c right)**.

To evaluate the replicability of the EC-GMR fabrication process, five different EC-GMRs with a 60 nm thick ITO layer were fabricated using identical processes. The optical characteristics of these EC-GMR sensors in terms of the resonance wavelength and Q factor are shown in supplementary table 1. The fabrication process consistently results in EC-GMR sensors that successfully support an optical resonance. The resonance wavelength for these five sensors was seen to vary by 5.2%. We note, the absolute resonance wavelength is not critical for the application of the EC-GMR as a biosensor as measurements are typically performed relative to the initial resonance wavelength.

FDTD simulations indicated that the optical resonance of the EC-GMR is influenced by the application of bias voltages, as has been observed in many combined electro-optical techniques^19^. To measure the effect of bias voltage on the EC-GMR, we monitored the resonance wavelength of a 60 nm thick ITO coated EC-GMR while applying increasing negative bias voltage to the ITO layer in 100 mM KPi buffer at pH 7.2 (**Fig. 3d**). Here, electrochemical measurements were performed in a three-electrode cell configuration using a Pt wire counter electrode and the ITO as the working electrode which was biased relative to a Ag/AgCl reference electrode.

The wavelength was seen to be stable up to a voltage of −0.6 V vs Ag/AgCl, after which the resonance wavelength was seen to shift to lower wavelengths up to a maximum applied voltage of −1 V vs Ag/ AgCl where the resonance wavelength had reduced by 0.35 nm. The ITO surface is negatively charged at pH 7.2 resulting in a Stern layer of predominantly positively-charged potassium ions at the ITO surface^32^ which will further increase when a negative bias is applied to the ITO. This high surface concentration of positively charged ions at the ITO-electrolyte interface will induce accumulation of electrons at the ITO surface. From the Drude model, the permittivity of the ITO is dependent on the free electron carrier density, such that the accumulation of electrons at the ITO surface will reduce the permittivity of the ITO and consequently lead to the observed negative resonance wavelength shift. The simulations of Fig. 2d predicted a linear relationship between resonance wavelength and the refractive index of the accumulation layer, however we observe a voltage window (between 0 and −0.6 V vs Ag/AgCl) where the resonance wavelength is stable. This may relate to the counteracting processes of bias driven depletion of electrons from the ITO and the accumulation of positive charge on the surface from ions in the media.

### Electrical and Optical Refractive Index Sensitivity

To quantify the refractive index sensitivity of the EC-GMR, a GMR sensor capped with a 60 nm thick ITO layer was integrated into a bespoke PDMS flow cell and exposed to solutions containing an increasing concentration of NaCl (0.1% to 20% w/v in ddH2O). The resonance wavelength increases with NaCl concentration, starting at 889.5 nm for 0.1% NaCl and increasing to 892.5 nm for 20% NaCl, reflecting the increasing refractive index of the solutions (**Fig. 4 a**). The bulk RI sensitivity was determined to be 84.4 nm/RIU a with a limit of detection (LOD) of 4.4× 10^−3^ RIU (**Fig. 4 b**). Previous GMR gratings have been reported which have a bulk sensitivity in the range of 84-140 nm/RIU^28^ . The relatively low optical performance is likely due to the reduced Q factor associated with optical scattering from the ITO layer. Improving the surface quality of the ITO and using improved fitting procedures such as Kalman filtering projection is likely to improve LOD in the future ^33^.

**Figure 4.**
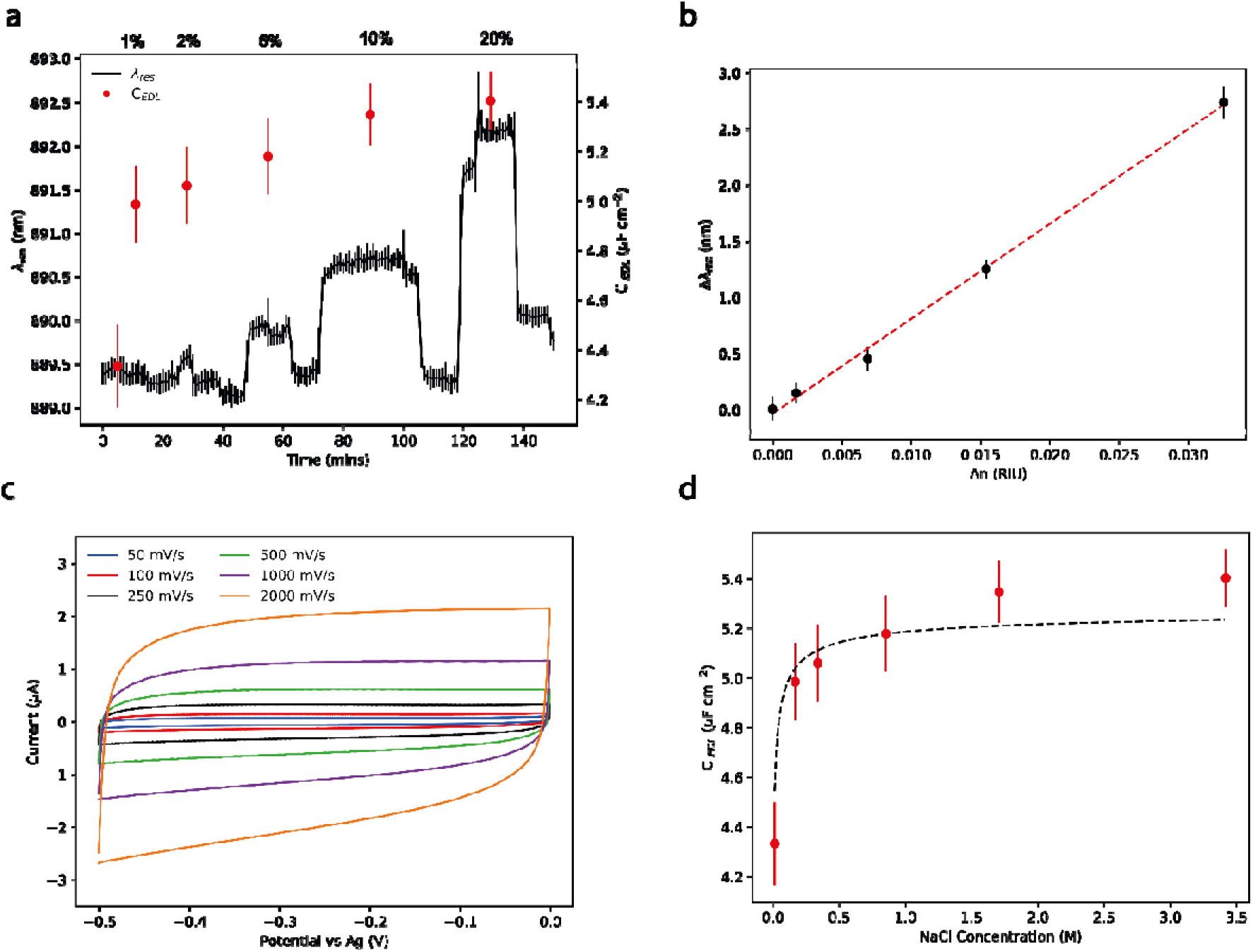
Simultaneous refractive index sensing and quantitative measurement of the electrochemical double layer: **a** Optical sensitivity measurement of the EC-GMR using increasing concentrations of 0.1, 1,2,5,10 and 20% NaCl in DI water with parallel measurement of double layer capacitance C_EDL_. Fitted resonance wavelength against time (black line). Red circles show the fitted double layer capacitance from CV scans taken at the same time. **b** Change in resonance wavelength Δλres with respect to change in NaCl refractive index Δn. Linear fit R 2= 0.997. **c** CV of EC-GMR with 0.1% NaCl solution at increasing scan rates. **d** Fitted C EDL against NaCl molarity, black dashed line is the simulated C_EDL_ assuming highly polarised water in the Stern layer ⍰ r = 6.

Cyclic voltammetry was performed simultaneously to quantify the electrochemical double layer capacitance. The voltage was scanned between 0 to −0.5 V vs Ag/AgCl over which range the bias induced shift in resonance for a EC-GMR coated with a 60 nm thick ITO layer was negligible (as shown in **Fig. 3d**). The absence of redox peaks over this voltage range and the symmetry between the anodic and cathodic currents indicate that we are measuring the non-Faradaic current associated with charging the double layer capacitance only rather than charge transport across the electrode-electrolyte interface (**Fig. 4 c**). The double layer capacitance, *C_EDL_*, at each salt concentration are shown in Figure 4a which were obtained from *i* = *C_EDL_v* where *v* is the voltage scan rate and *i* is the current at the centre of the scan range. The measured values of *C_EDL_*were compared to theoretical calculations based on a serial capacitor model of the electrochemical double layer (Eq. 1) where *C_dl_*and *C_sl_*are the diffuse and Stern layer capacitance respectively and are given by the parallel plate capacitor equation (Eq. 2) where A is the electrode area, ε_0_ is the permittivity of vacuum, ε_r_ is the relative permittivity and where is layer thickness.

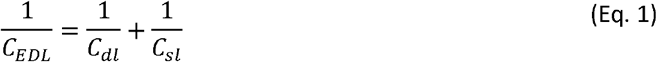

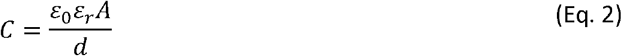

The thickness of *C_dl_*is determined by the Debye length, κ^-1^, for a monovalent salt, where F is Faraday’s constant, R is the gas constant and C_0_ is the salt concentration in mol/l (Eq.3).

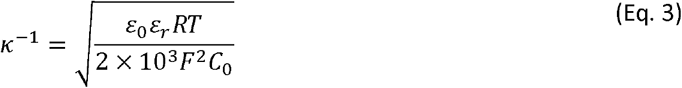

The thickness of *C_sl_*is 6 Å, estimated from the size of hydrated Na^+^ and Cl^−^ ions. The water molecules in the Stern layer are highly polarised under the high electric field (833 MV/m at maximum voltage) so a reduced water permittivity of ε_r_ = 6 is used^34^. The combined model accounts for the domination of the Stern layer capacitance at higher NaCl concentrations. The measured values of *C_EDL_* for each NaCl concentration are all within 6% of the theoretical *C_EDL_* when using the reduced permittivity for the Stern layer (**Fig. 4 d**). The comparison of the optical and electrochemical response also shows that *C_EDL_* is dominated by the Stern layer of adsorbed ions onto the electrode surface as *C_EDL_* remains largely constant in comparison to the refractive index change of the media with increasing NaCl concentration.

We have demonstrated that the EC-GMR is capable of performing quantitative optical and electrochemical measurements simultaneously. The optical sensitivity of the EC-GMR is reduced compared to previously reported GMR sensors, due to optical losses in the ITO layer. This loss in sensitivity is however offset by the gain of the entirely new electrochemical modality which has not been reported before using GMR structures.

### Decoupling surface adsorption and quasi electrochemical behaviour of redox active molecules by multimodal sensing

Finally, we demonstrate the ability of the EC-GMR to characterise both the refractive index and electrochemical properties of a solution of redox active molecules in parallel. We focus here on methylene blue (MB), a pH sensitive redox active molecule that reduces to leucomethylene blue in a 2 electron 1 proton process^35^ and has become a widely used redox reporter in applied and basic biomedical research, including microRNA detection^36^ DNA damage reporting^37^ antibiotic susceptibility^38^, eukaryotic cell viability^39^ and aptamer antigen detection assays^40^. MB is an attractive redox active reporter because of its chemical stability, electrochemical reversibility, pH sensitivity and unambiguous redox potential compared to the electrochemical background observed on typical electrode materials^41^. However, the electron transfer process is dependent on buffer properties, such as pH,^26^ surface chemistry and on the voltametric techniques themselves which can regulate the mechanism of electron transfer^42^. It thus remains challenging to deconvolute each of these contributions using electrochemical measurements alone. Here we show how greater insight into the redox activity of MB can be provided by combining the complementary information revealed by the EC-GMR multimodal sensor.

The EC-GMR reflectance spectra show an increase in optical reflectance with concentration of MB due to the higher reflectance of MB solutions in the near infrared (**Fig. 5a**). We also observed an increase in resonance wavelength with increasing concentrations of MB corresponding to the increase in solution refractive index. The fitted resonance wavelength increased exponentially with increasing concentrations of MB, increasing by 0.45 nm between the KPi electrolyte and 1 mM MB in KPi (**Fig. 5b**). The LoD of the EC-GMR for the detection of MB is 60 µM, showing a slight improvement over existing evanescent field methods for the detection of redox molecules such as MB and with a larger dynamic range ^43^.

**Figure 5.**
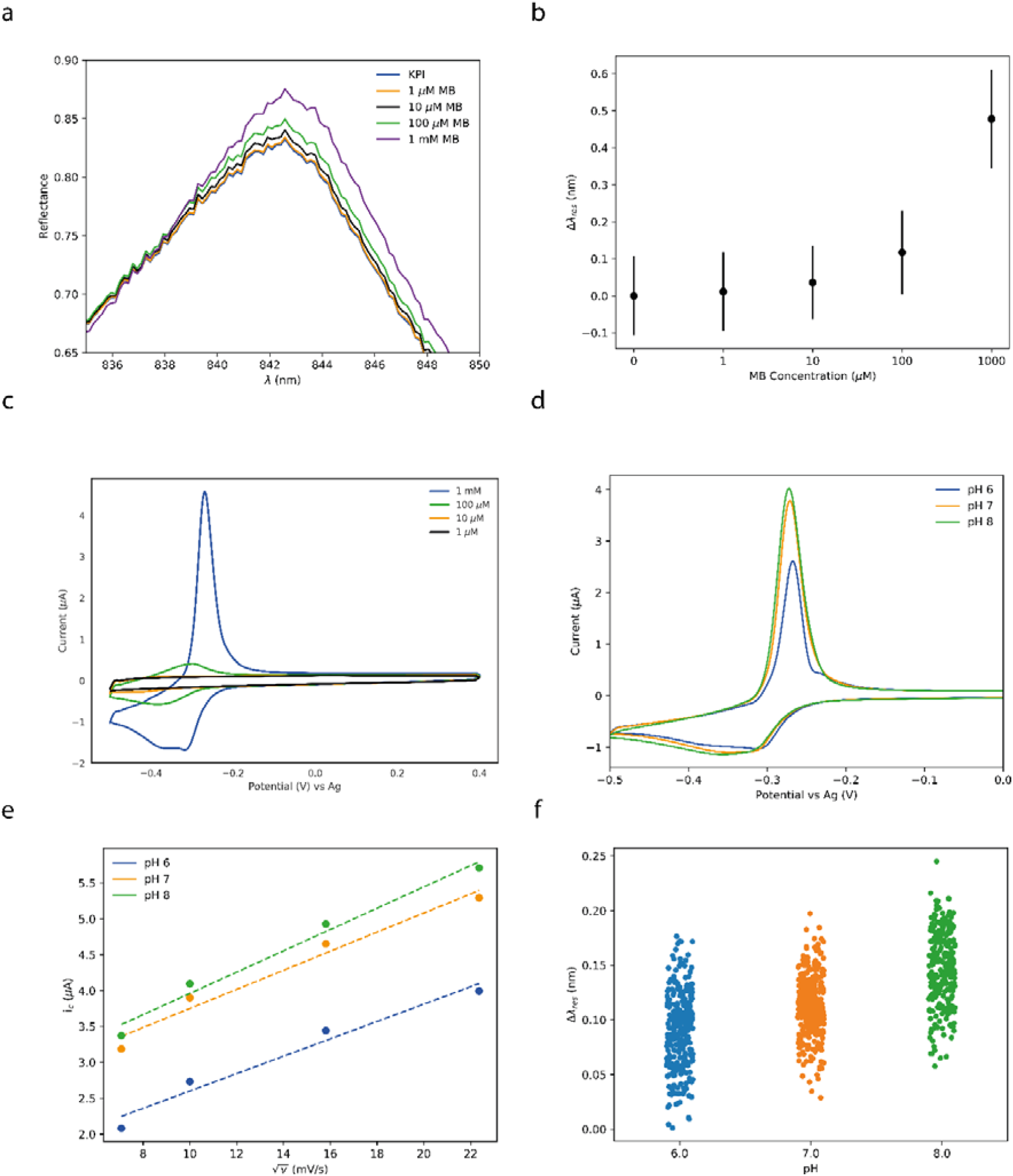
Optical and electrochemical characterisation of methylene blue in solution: **a** Reflectance spectra of EC-GMR in KPi buffer and MB concentrations from 1 µM to 1 mM. **b** Mean change in resonance wavelength from KPi with increasing concentration of MB from 1 µM to 1 mM in 10 mM KPi buffer. Resonance wavelength was recorded while the CV measurements were taking place. Error bars are standard deviation (n = 100). **c** CV plot of MB concentrations from 1µM to 1 mM in 10 mM KPi buffer using EC-GMR as the working electrode. **d** Cyclic voltamagrams of 100 µM MB in pH 6, 7, and 8. **e** MB peak reduction current against the square root of the scan rate for different Ph. Dashed lines indicate a linear trend. **f** Change in EC-GMR resonance wavelength against MB concentration at pH 6-8 normalised to the 0 MB.

CV measurements at each MB concentration from 1 µM to 1 mM were performed simultaneously with the optical measurements, demonstrating that the EC-GMR is able to support refractive index sensing in parallel with Faradaic processes as required for electrochemical measurements of redox active molecules (**Fig. 5 c).** While the redox peaks can be determined for MB concentration between 1 mM and 100 µM using CV (see Supplementary Table 2), the large, non-Faradaic background current masks the reduction and oxidation peaks at lower concentrations. The anodic and cathodic redox potentials, *E_pa_* and *E_pc_* respectively, show good agreement with previously published values for MB in phosphate buffer using an Ag/AgCl reference electrode ^44^. The shift in peak potential and peak width with increasing MB concentration is likely due to the change in the buffer, such as pH and conductivity, that occurs upon addition of high concentrations of MB.

From Fig. 5c, we find that the separation between oxidation and reduction peaks for MB is 49 mV and 81 mV at 1 mM and 100 µM respectively (Supplementary Table 2). This is greater than the theoretical ideal peak separation expected for a two-electron redox reaction of 29 mV. Differences in the peak separation values are typically attributed to either the resistance of the working electrode limiting electron transfer ^45^ or physisorption of MB on to the ITO surface^46^. Moreover, as the redox reaction for MB is reversible, the peak reduction current, *I<sub>c</sub>* is expected to increase linearly with voltage scan rate. In contrast, here we observed that at low scan rates, the relationship is non-linear. This known phenomenon is attributed to either a quasi-reversible electrochemical reaction or to surface adsorption on the electrode, Critically, the explanation for the observed deviations from the theoretical ideal cannot be reconciled from the electrochemical measurements alone (a more detailed explanation is provided in the supplementary information).

To explore if the combined optical and electrical measurements can reconcile the observed differences from the ideal redox behaviour of the MB system, we measured the λ_res_ change and the corresponding CV profile of 100 µM MB in the pH range 6-8. As the MB redox reaction is a 2 electron, 1 proton process, the resulting voltammograms are expected to shift with pH, as can be clearly observed by the increase in redox potential with respect to increasing pH (**Fig. 5d, Supplementary Table 3**). We observe non linearity with respect to root of the scan rate for all observed pH (**Fig. 5e**) However, the charge of the MB will also change over this pH range, becoming increasingly less positively charged until around pH 8 when the MB will become charge neutral and thus hydrophobic, leading to greater physisorption on the EC-GMR sensor surface^47^. Evidence of MB physisorption is found in the shift in λ_res_ (normalised to the resonance wavelength of the electrolyte) which increases with pH (**Fig. 5f**), and is indicative of an increase in local refractive index that would be expected due to surface adsorption of MB. The ability to decouple surface adsorption from the electrochemical behaviour of the MB demonstrates clearly the complementary information that can be obtained from a multimodal measurement as the non-linearity can be attributed to apparent quasi electrochemical behaviour of the molecule, likely due to non-ideal electron transfer between the molecule and the electrode. We note, the insensitivity of the resonance wavelength to the solution pH does open opportunities of using redox active labels as pH sensors while performing simultaneous refractive index measurements.

## Conclusion

In this study we have presented the EC-GMR, a guided mode resonance biosensor with an ITO electrode capable of performing simultaneous optical and electrochemical measurements. We have shown that guided mode resonances can be theoretically supported in structures containing optical lossy materials such as transparent conductive oxides and that the resonance can be observed experimentally for the TE mode. The resonance wavelength is stable below −0.6 V of applied bias for a 60 nm ITO film which affords independent optical and electrochemical measurements. We have characterised the optical and electrical sensitivity and shown that while the optical sensitivity is lower than previously reported GMR structures we can also perform quantitative measurements of the electrochemical double layer within 10% of the expected theoretical predictions. Finally, we have demonstrated that the EC-GMR can be used for monitoring redox processes while performing optical measurements giving rise to the potential for simultaneous measurements of redox active reporter molecules in parallel with optical surface binding measurements. We believe that the parallel combination of quantitative electrochemical measurements and optical sensing using the robust low cost GMR platform will open novel opportunities for the interrogation of complex biological systems such as monitoring of bacterial biofilm adhesion combined with detection of redox quorum sensing molecules or enzymatic degradation of antibiotics by drug resistant bacteria.

## Funding Information

This research was supported by the Wellcome Trust four-year PhD programme (WT095024MA) ‘Combating infectious disease: computational approaches in translation science’ and an Engineering and Physical Sciences Research Council Programme Grant (EP/P030017/1).

## Author Contribution

**S.T:** Conceptualization; investigation; formal analysis; writing – original draft preparation**. S.B:** Investigation; validation writing – original draft preparation**. G.P:** Investigation; methodology; writing – review & editing. **C.K:** Investigation; writing – review & editing **C.P.R:** Validation; writing – review & editing. **T.F.K:** Conceptualization; writing – review & editing; supervision. **S.D.J:** Conceptualization; writing – review & editing; supervision.

## Methods

### Design and Simulation

The numerical modelling of the nanostructures was performed using a Finite difference time domain (FDTD) method with commercially available FDTD software (Lumerical FDTD Solutions). The 1D periodic structure was modelled as a 150 nm thick Si_3_N_4_ grating, with a period of 555 nm and a 0.8 duty cycle. The Si_3_N_4_ layer was assumed to have a uniform refractive index of 1.89 with the loss factor assumed to be negligible. The ITO layer was modelled as an continuous layer, with a uniform thicknesses on the top and in the troughs of the grating, whilst the thickness of the ITO on the sidewalls was varied as a percentage of the total ITO thickness. The photonic and electrical properties of the ITO layer were extracted from previous literature^48^. The accumulation layer within the ITO slab was modelled as a 1 nm region of differing refractive index located at the ITO-electrolyte gate interface. The simulations were performed assuming illumination with a spatially coherent plane wave and periodic boundary conditions were applied on all sides of the computational domain. For accumulation layer simulations, a minimum mesh step of dz = 0.5 nm was used, so as to accurately resolve the layer.

### GMR Fabrication

150 nm Si3N4 wafer on borosilicate glass (Silson) was diced into 15 x 15 mm squares (Disco DAD-321 wafer saw, Giorgio Technology), cleaned by sonication in acetone and iso-propyl alcohol (IPA) then dried under nitrogen gas. A layer of positive electron beam resist (AR-P 6200.13, All Resist) was spin coated onto the wafer at 5000 RPM then baked at 180 ⍰C for 7 minutes. A charge dissipation layer (AR-PC 5090, All Resist) was next spin coated at 2000 RPM then baked at 100 ⍰C for 2 minutes. The GMR structure was patterned using a Raith Voyager electron beam lithography system using a base dose of 130 µC and a dose factor of 1.1. After exposure, the AR-PC was removed by immersing for 2 minutes in room temperature DI water then dried under nitrogen. The AR-P 13 was developed for 2 minutes in xylene, the excess xylene removed by dipping into DI water, followed by IPA, before drying under nitrogen. The GMR pattern was etched into the Si3N4 substrate using reactive ion etching (RIE) with a (29:1 ratio of CHF3: O_2_) at a base pressure of 1.9 × 10^−1^ mbar for 7 minutes. The remaining resist was removed by sonication for 2 minutes in micropositTM remover 1165 (MicroChem) followed by acetone then IPA.

### ITO Deposition

ITO was deposited using an electron beam evaporation system (MBraun) with In_2_O_3_SnO_2_ evaporation pellets (90:10 % Testbourne) starting with a base pressure of 10^−6^ mbar before flowing in 7.5 SCCM O_2_ to give an evaporation pressure 10^−4^ mbar. The ITO was deposited at 0.5 Å/s after two rise and soak steps to remove impurities from ITO using an 8 kV beam voltage. To improve optical transmission and conductivity, the structure was annealed for 1 hour at 4001C in an O_2_ atmosphere. After annealing, a wire was bonded onto the ITO using silver conductive epoxy resin mixed 1:1. The resin was then baked at 90 ⍰C until cured.

### Optical and Electrochemical Sensitivity measurements

To fabricate the channel master, SU-8 was spin coated onto an Si wafer then exposed using electron beam lithography. After development the sample was hardbaked for 15 minutes at 150 1C. An antiadhesion layer was formed by placing the master in a vacuum chamber alongside a drop of Trimethylchlorosilane on a petri dish for 15 minutes. The channel dimensions were 1 mm wide, 10mm long and 100 µm high. The polydimethylsiloxane (PDMS) channel was made by mixing the silicon elastomer and curing agent at a 10:1 ratio. The PDMS was poured onto the mould and then degassed in a vacuum chamber for 15 minutes until all of the bubbles in the mix had been removed. The PDMS was baked at 60 ⍰C overnight then removed from the mould, cut to size using a scalpel and then hole punched to allow access to the fluidic tubing. To adhere the PDMS channel to the ITO surface, the sensor and the channel were treated with O_2_ plasma (22 SCCM of O_2_ for 25 seconds at DC voltage of 180V). The PDMS was then aligned to the GMR substrate and the assembly was baked at 60 ⍰C overnight to permanently seal the PDMS to the sensor surface. After bonding, tygon tubes were inserted into the holes punched in the PDMS. Samples were flushed through the channel using a syringe pump operating at 20 µl min^−1^. For electrochemical measurements, the Pt counter electrode and Ag pseudo reference were placed in the waste receptacle for the PDMS flow cell drain tube. The limit of detection (LOD) was determined as 3 σ, where σ is the standard deviation of the system noise which was determined by monitoring the resonance wavelength at a fixed illumination for 500 s.

Potassium phosphate (KPi) buffer was prepared by mixing 71.7 mL of 1 M K_2_HPO_4_ with 28.3 mL 1 M KH_2_PO_4_ dissolved in DI water. The 1 M buffer was diluted 1:10 with DI to obtain a pH 7.2 100 mM KPi buffer. The pH was validated using a pH meter (Mettler Toledo). Methylene blue (Sigma) was diluted with 100 mM KPi buffer. Solutions of increasing NaCl concentration were prepared with DI water and NaCl (Sigma). The refractive index of the NaCl solutions was measured using a refractometer (name, supplier).

Electrochemical measurements were performed using an SP200 potentionstat (Bio logic) using a Pt wire as a counter electrode and either a Ag/AgCl reference or a Ag pseudo reference electrode.

The resonance wavelength was measured in reflection using a custom optical setup (Fig 1). An HL 2000 tungsten halogen light source (Ocean Optics) is focused into the back focal plane of a NeoDplan 10x (NA 0.25) objective to produce a collimated incident light source. The reflected light from the EC-GMR is coupled into a fibre and the spectra measured using a CCD spectrometer meter (Thorlabs). A CMOS camera Coolsnap Dyno Photometrics) is used for focusing and alignment of the GMR sensors. Prior to each measurement, the GMR reflectance was normalised to the Ag coated mirror reflectance spectra. A custom Labview script was used for recording the reflectance spectra over time. The measured spectra was fitted to a Fano resonance in Python using the *numpy* package to extract the resonance wavelength. Hyperspectral imaging was performed using the same optical setup but using a halogen white source (Spectral Products) coupled to a CM110 1/8 monochromator (Spectral Products). Brightfield images were taken by the camera at each illumination wavelength from 800 - 900 nm at 0.5nm steps. The hyperspectral image is made by selecting the highest reflectance wavelength for each pixel in the image.

### Methylene Blue Redox Sensing

Concentrations of methylene blue from 1 µM to 1 mM diluted in 100 mM KPi buffer were pipetted on to the EC-GMR sensor in a custom PDMS well adhered to the sensor surface. Using a static well as opposed to the microfluidic flow cell allows the reference electrode to be placed closer to the ITO working electrode, reducing the Ohmic drop between the working and reference electrode. The sensor was washed three times with KPi buffer between each concentration to remove MB and salts bound non-specifically to the surface. The reflectance was monitored every two seconds and the resonance wavelength was extracted by fitting the measured reflectance to the Fano equation. The redox activity of the MB was probed by CV and square wave voltammetry using an SP200 potentiostat and a Pt wire counter electrode and an Ag wire pseudo reference electrode. The MB concentration was limited to 1 mM by the solubility of the MB; higher concentrations were found to precipitate out of solution at the tested pHs.

## Notes

### Competing Interest Statement

The authors have declared no competing interest.

