## Supplementary Information for "Electrochemical guided mode resonance biosensor for simultaneous refractive index and electrochemical measurements"

**Interpretation of Cyclic Voltammetry**

Voltammograms are very powerful tools for interrogating redox active species, but the complete electrochemical behaviour is dependent on many parameters such as the buffer solution, electrode material, analyte concentration, analyte diffusion, temperature, and voltammogram scan rate. It is also important to note that the electrochemical response is strongly influenced both by interactions of the redox active molecules with the surface of the electrode and their behaviour in free solution.

The peak current, I_p_, of a freely diffusing redox active molecule from a CV voltammogram is given by the Randles-Sevcik equation (Eq S1) where *n* is the number of electrons in the redox reaction, *F* is Faraday’s constant, *v* is the scan rate, *D* is the diffusion coefficient, *R* is the gas constant, *T* is the temperature and *C* is the concentration. The diffusion coefficient, *D*, is also directly affected by changes in pH, which we show in Table 4, therefore the differences in *I_c_* with scan rate for different pH cannot be attributed to surface adsorption alone.

$$i_{p}=0.4463nFAC\sqrt{\frac{nFvD}{RT}}$$

(Eq S1)

From Eq S1, the peak current on a CV plot should scale linearly with the square root of the scan rate for a freely diffusing molecule, but we observe a deviation from the linearity (Fig 5f). For a surface immobilised molecule, the peak current scales linearly with the scan rate which we also do not observe (Fig S3) but this deviation could also be due to limitations in the electron transfer to the electrode^1^.

The peak separation, *E_pc_ - E_pa_*, from the CV can also be used to determine if there is surface adsorption. An ideal surface analyte will have no separation between *E_pc_* and *E_pa_*. A completely reversible reaction of a freely diffusing molecule will have a *E_pc_ - E_pa_* separation of 57 mV/*n* where *n* is the number of electrons in the reaction. We first observe that none of the measurements match the theoretical value and a large charge in peak separation at pH 7 with scan rate (Fig S4a). On further inspection of E_pc_ and E_pa_ we observe that the E_pc_ remains constant and that E_pa_ changes with scan rate but given the broad oxidation peak it is difficult to define E_pa_ accurately (Fig S4b). Quasi reversible reactions show an increase from the ideal scaling with scan rate, however this can also be attributed to limitation of electron transfer by non-ideal electrodes, such as if the electrode were to become fouled.

It is possible to also to attempt to subtract some of these features from the voltammograms by performing analyte free measurements, to remove the non-faradaic contribution to the CV. This approach is challenging however as the analytes themselves can induce changes in the non-faradaic contribution and therefore also requires careful interpretation^2^.

There is therefore no completely unambiguous way to know from the electrochemical measurements alone that pH dependent surface adsorption is taking place. The refractive index sensing however is only sensitive to the local refractive index at the surface once normalisation for buffer pH has been performed, allowing surface adsorption to be confirmed unambiguously.

To summarise:

- I_c_ depends on scan rate and diffusion coefficient, surface effects included adsorption are included in this.
- Deviation from linearity in I_c_ with scan rate can depend on surface adsorption and electrode electron transfer kinetics.
- Changes in peak potential separation are due to both surface adsorption and electrode electron transfer kinetics.
- Refractive index measurements only due to presence of different refractive index material at the surface.

**Figures**

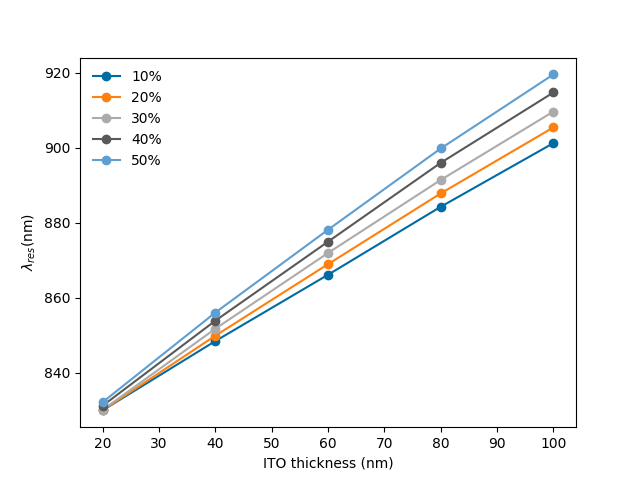

**Supplementary Figure 1**: FDTD simulations of the EC-GMR grating with increasing sidewall thickness of ITO as a percentage of the ITO thickness at the top and bottom of the grating.

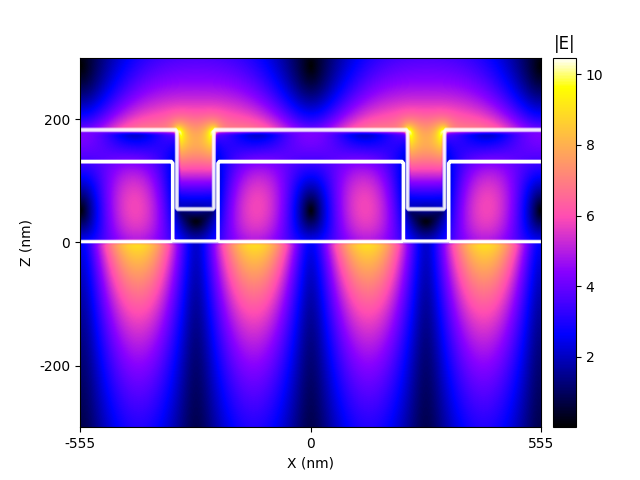

**Supplementary** **Figure 2**: Electric field profile of the TM mode at λ_res_ for the EC-GMR coated in 60 nm ITO with 40% sidewall infill.

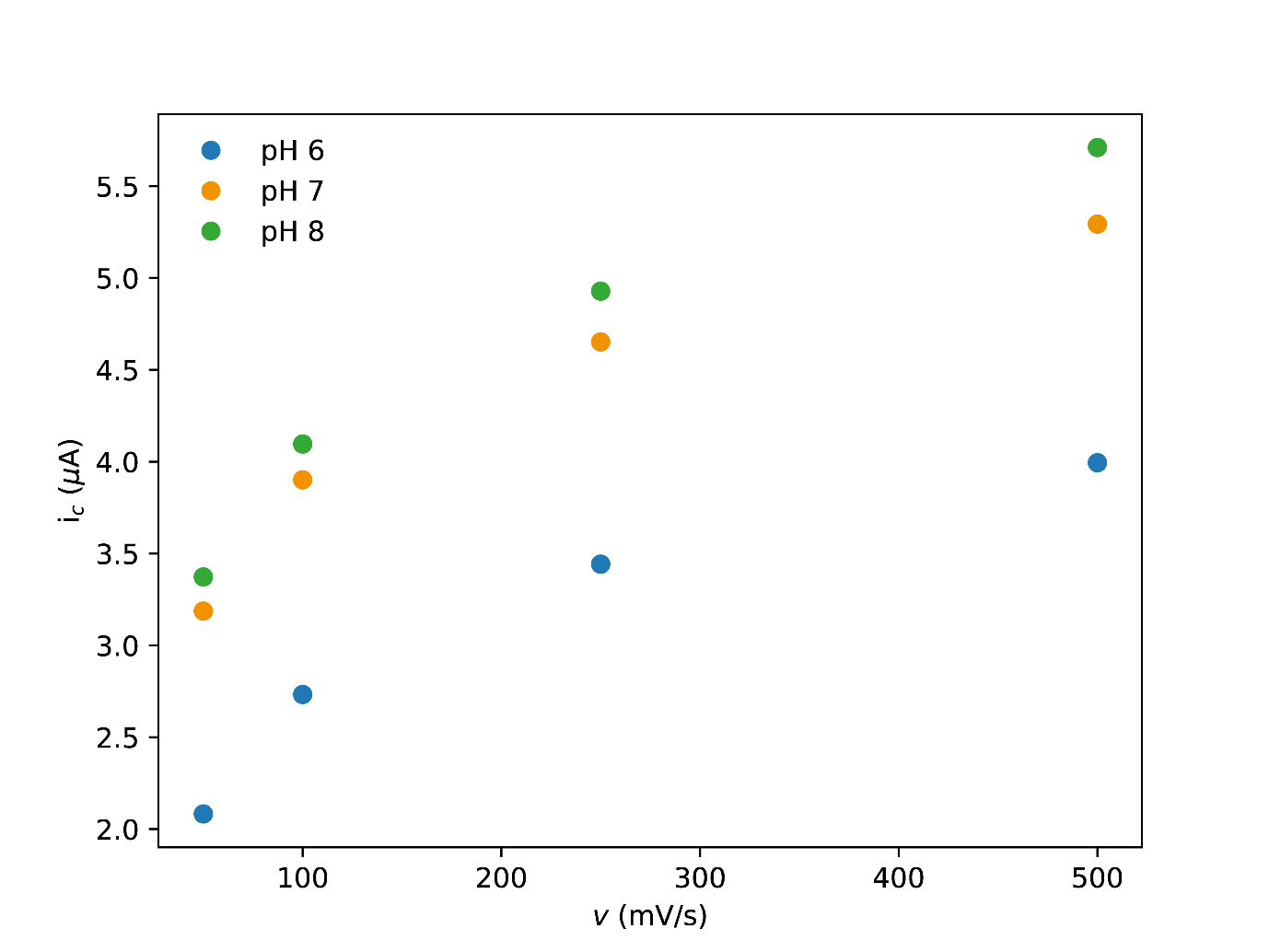

**Supplementary** **Figure 3**: Peak cathodic current of MB between pH 6 – 8 recorded using an EC-GMR with a 60 nm thick ITO electrode as a function of cyclic voltammetry scan rate, *v*.

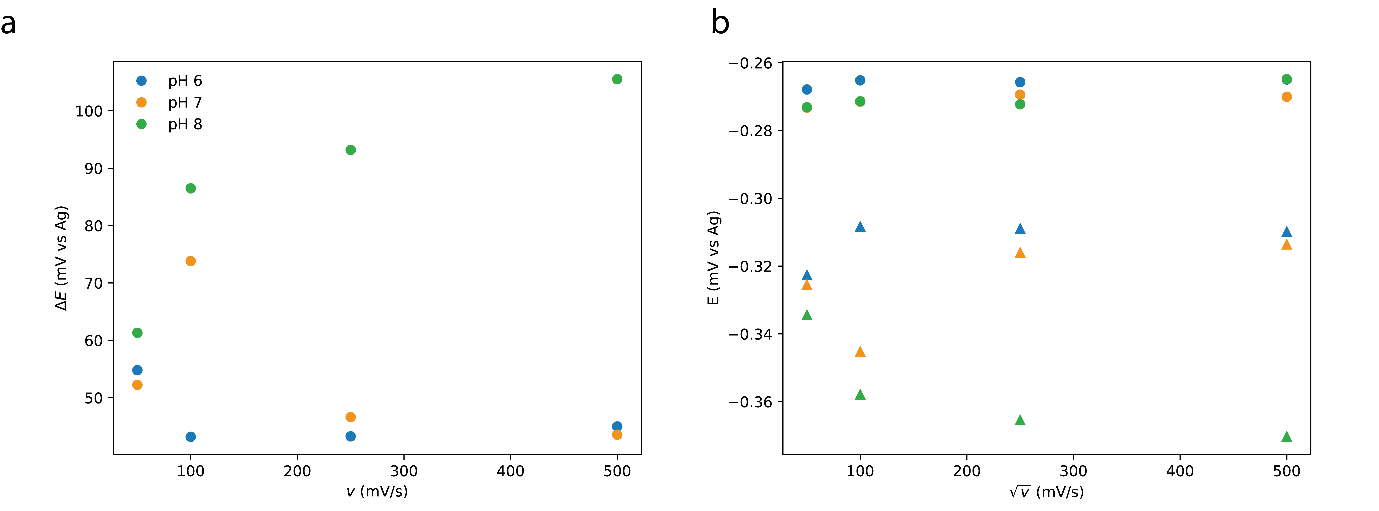

**Supplementary** **Figure 4**: a) peak separation and b) E_pa_ (circles) and E_pc_ (triangles) of MB between pH 6 – 8 recorded using an EC-GMR with a 60 nm thick ITO electrode as a function of cyclic voltammetry scan rate, *v*.

**Tables**

Table 1: Measured resonance parameters of Si_3_N_4_ GMR, 20 nm and 60 nm ITO EC-GMR sensors

| Sample | λ_res_ (nm) | FWHM (nm) | Q factor |
| --- | --- | --- | --- |
| Si_3_N_4_ | 830 | 9.14 | 90 |
| 20 nm ITO | 854 | 10.42 | 82 |
| 60 nm ITO | 882 | 12.5 | 70 |

Table 2: Measured parameters of 60 nm ITO EC-GMR fabricated independently

| Sample | λ_res_ (nm) | FWHM (nm) | Q factor |
| --- | --- | --- | --- |
| 1 | 852 | 9.6 | 88.76 |
| 2 | 882 | 11.28 | 78.19 |
| 3 | 855 | 9.15 | 89.88 |
| 4 | 842 | 11.0 | 76.57 |
| 5 | 868 | 11.72 | 74.05 |

Table 3: Parameters of MB extracted from CV measurements. CV peaks for 10 *µ*M and 1 *µ*M could not determined.

| MB Concentration (µM) | $E_{pa}$ (mV) | $E_{pc}$ (mV) | Peak Separation (mV) |
| --- | --- | --- | --- |
| 1000 | -317 | -268 | 49 |
| 100 | -376 | -295 | 81 |

Table 4: Electrochemical parameters of MB at pH 6-8.

| pH | $E_{pa}$ (mV) | $E_{pc}$ (mV) | Peak Separation (mV) | *D (*10^-6^ cm^2^s^-1^) |
| --- | --- | --- | --- | --- |
| 6 | -267 | -307 | 40 | 1.422 |
| 7 | -271 | -314 | 43 | 1.717 |
| 8 | -273 | -321 | 49 | 2.139 |
